# Live imaging reveals polarized calcium transients during plant pathogen development and host colonization

**DOI:** 10.64898/2026.05.14.725065

**Authors:** Max HJ Pluis, Djenatte Abdennour, John J. Mackrill, Edouard Evangelisti

## Abstract

Ca^2+^ signaling mediates rapid cellular responses across eukaryotes, but its spatiotemporal dynamics remain largely inaccessible in many genetically less tractable microbial lineages. Oomycete plant pathogens, including *Phytophthora* species, undergo rapid transitions between motile, encysted, germinating, and invasive stages, yet the organization of Ca^2+^ dynamics during these transitions is poorly understood. Here, we adapt the genetically encoded ratiometric biosensor MatryoshCaMP8s for *in vivo* calcium imaging in *Phytophthora palmivora*. The reporter is stably expressed without major detectable effects on sporulation or virulence and reports rapid ratiometric responses to cold shock and calcimycin/A23187. Using live-cell imaging, we uncover stage-specific Ca^2+^ dynamics across the pre-infective and early infection cycle. Sporangia approaching zoospore release display spatially heterogeneous Ca^2+^ transients, newly formed cysts occasionally exhibit Ca^2+^ transients, and germinating cysts show recurrent Ca^2+^ transients at the germ tube tip. Similar sharp ratiometric pulses occur during early plant infection, indicating that polarized Ca^2+^ transients are not restricted to *in vitro* germination but recur at the host surface. Together, our work establishes live ratiometric calcium imaging in oomycetes, reveals polarized Ca^2+^ transients as recurrent signatures of developmental transitions and early host colonization, and opens the way to mechanistic dissection of signaling, polarity, and infection in a major group of plant pathogens.

## Introduction

Oomycetes are filamentous eukaryotic microbes that include some of the most destructive plant pathogens affecting crops and natural ecosystems. The genus *Phytophthora*, for example, comprises more than 200 species, most of which are associated with plant disease [1]. Notably, *Phytophthora infestans*, the causal agent of potato late blight, triggered the Great Irish Famine and continues to cause billions of dollars in losses annually [2,3]. The tropical species *Phytophthora palmivora,* which is the focus of this study, infects a wide range of staple and cash crops, including cacao, oil palm, mango, pineapple, rubber, and citrus [4]. *Phytophthora* pathogens rely on motile biflagellate zoospores to navigate their environment, respond to chemical and ionic cues, and reach suitable infection sites at the plant surface [5,6]. Upon reaching these sites, zoospores shed their flagella and build a cell wall, forming cysts. Cysts then germinate and enter host tissues by developing specialized penetration structures called naifu appressoria [7,8]. These appressoria rely on localized reorganization of the actin cytoskeleton to sharpen the invasive hyphal tip, thereby acting as microscopic knives that slice through the cuticle and plant cell wall rather than relying primarily on high turgor pressure [9,10]. These rapid transitions from motility to adhesion, polarity, and invasion imply tight intracellular coordination, yet the signaling dynamics underlying them remain largely unknown.

Ca^2+^ acts as a universal second messenger in eukaryotic cells, transducing environmental and developmental cues into rapid cellular responses [11,12]. Transient changes in cytosolic Ca^2+^ are generated through influx from the extracellular space or release from intracellular stores, shaped by Ca^2+^ transporters and buffers [13,14], and decoded by a broad range of Ca^2+^-binding and Ca^2+^-regulated proteins. These notably include EF-hand-containing regulators such as calmodulins, as well as kinases, phosphatases, and motor proteins [15–17]. In plant-microbe interactions, Ca^2+^ transients shape both immunity and symbiosis. For instance, pathogen perception and herbivore attack trigger rapid cytosolic Ca^2+^ elevations that contribute to immune activation, whereas symbiotic interactions with rhizobia or arbuscular mycorrhizal fungi induce nuclear Ca^2+^ spiking decoded by Ca^2+^/calmodulin-dependent kinases to activate symbiotic gene expression [16,18,19]. Monitoring these dynamics in living cells has long relied largely on small-molecule fluorescent calcium indicators, such as the BAPTA-derived dyes Fura-2 and Fluo-4 [20]. However, such approaches depend on efficient dye delivery, retention, and intracellular activation, can perturb cells, and often lack the temporal or spatial resolution required to track rapid subcellular events [21]. Genetically encoded calcium indicators, such as the GCaMP family [22], overcome some of these limitations by linking Ca^2+^ binding to changes in fluorescence. In filamentous fungi, expression of the FRET-based Cameleon sensor previously revealed pulsatile Ca^2+^ signatures associated with hyphal growth and selected infection-related contexts [23]. Ratiometric designs, including MatryoshCaMP [24], further incorporate a Ca^2+^-insensitive reference fluorophore, enabling more robust measurements by correcting for variation in expression level, cell geometry, and image acquisition.

In oomycetes, changes in cytosolic Ca^2+^ concentration have been implicated in several key processes of the infection cycle, including sporangial discharge [25], zoospore swimming behavior [26], chemotaxis [27], and hyphal growth [28,29]. Pioneering work used microinjected Fura-2 dextran to show that cold treatment induces rapid Ca^2+^ elevations in *Phytophthora* sporangia [30]. More recent studies used dye-based approaches to provide evidence for Ca^2+^ signaling downstream of receptor-like kinase-mediated sterol perception [31]. However, because genetic manipulation of oomycetes remains challenging, live genetically encoded calcium imaging has not been established in these organisms, leaving the spatiotemporal organization of Ca^2+^ signaling largely inaccessible. As a result, we still do not know whether Ca^2+^ acts as a diffuse cellular signal or as a localized, stage-specific regulator during the developmental transitions that precede host penetration and accompany early infection.

Here, we adapted the ratiometric MatryoshCaMP design to *Phytophthora palmivora* and established a stable reporter line for *in vivo* calcium imaging. We validated the biosensor using cold shock and the ionophore calcimycin/A23187, then used it to monitor Ca^2+^ dynamics across successive pre-infective and early infection stages. We reveal spatially heterogeneous Ca^2+^ transients in sporangia prior to zoospore release, rare Ca^2+^ transients shortly after encystment, recurrent tip-associated Ca^2+^ transients and Ca^2+^ waves propagating along germ tubes during cyst germination, and sharp Ca^2+^ pulses during early plant infection. Together, our work establishes live ratiometric calcium imaging in oomycetes and reveals previously inaccessible polarized Ca^2+^ transients as recurrent signatures of developmental transitions and early host colonization.

## Results

### A ratiometric MatryoshCaMP8s reporter can be stably expressed in *Phytophthora palmivora*

Ratiometric biosensors provide an internal fluorescence reference that enables the quantification of molecular dynamics in living cells despite variations in expression level, cell geometry, or imaging conditions [32]. To establish such a system for calcium imaging in *Phytophthora*, we generated the pHygro-MatryoshCaMP8s vector. This construct combines a hygromycin resistance cassette, chosen to maintain compatibility with overexpression and CRISPR/Cas plasmids conferring geneticin (G418) resistance, with an expression cassette encoding a MatryoshCaMP calcium reporter [24] (**Fig. 1a**). The reporter retains the sensory module and engineered mutations of jGCaMP8s [22]. It also contains an mScarlet-I fluorescent protein carrying the E95D mutation (corresponding to E220D in the full MatryoshCaMP8s reporter sequence) to reduce puncta formation [33], inserted between flanking GGT/GGS residues and serving as an internal reference fluorophore for ratiometric measurements (**Fig. 1b**). Transformation of *P. palmivora* yielded seven independent transformant lines. Among them, lines #6 and #7 showed colony morphologies comparable to the wild type, whereas line #3 displayed severely reduced growth and was excluded (**Fig. 1c**). A slight delay in radial growth was observed on selective medium. Both selected lines showed sporulation levels comparable to the wild type (**Fig. 1d**). However, line #6 displayed a homogeneous mScarlet-I signal in axenic culture, consistent with stable reporter expression, whereas sectors lacking a detectable mScarlet-I signal were observed in line #7 (**Fig. 1e**). We therefore focused subsequent analyses primarily on line #6. We further assessed the virulence of line #6 on *Nicotiana benthamiana* leaves (**Fig. 1f**). No significant difference in pathogenicity was observed between line #6 and the wild type. Together, these results identify line #6 as a suitable reporter line for live ratiometric calcium imaging in *P. palmivora*, with robust expression and no major detectable impact on sporulation or virulence.

**Figure 1.**
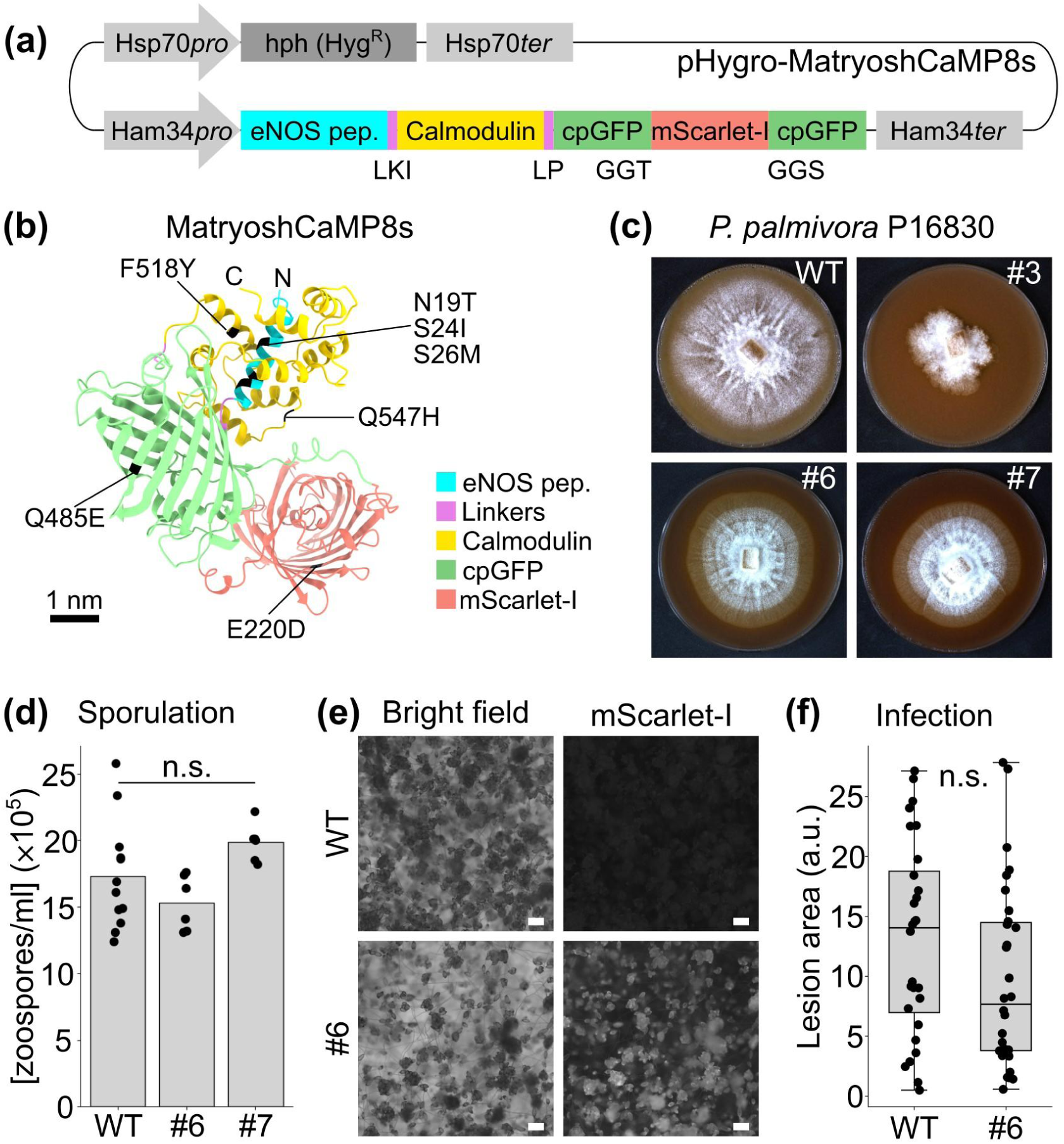
Development of a genetically encoded ratiometric calcium biosensor in *Phytophthora palmivora*. **(a)** Schematic representation of the pHygro-MatryoshCaMP8s construct. The vector carries a hygromycin resistance cassette (Hyg^R^) driven by the Hsp70 promoter, and an expression cassette under the Ham34 promoter to express the MatryoshCaMP8s calcium biosensor. The reporter consists of an eNOS-derived peptide, calmodulin, a circularly permuted GFP (cpGFP), and an mScarlet-I fluorescent protein inserted in a permissive loop of the GFP. **(b)** AlphaFold 3 model of MatryoshCaMP8s, showing the eNOS peptide, linker regions, calmodulin, cpGFP, and mScarlet-I modules. Mutations introduced in the sensor are highlighted in black. Color code as indicated. **(c)** Colony morphology of *P. palmivora* wild type (WT) and representative MatryoshCaMP8s transformant lines (#3, #6, and #7) grown on selective medium. **(d)** Quantification of zoospore production in WT and transformant lines #6 and #7. Bars represent mean values, and dots indicate individual measurements. No significant differences were detected between genotypes (Kruskal-Wallis test, P= 0.26). **(e)** Bright-field and RFP fluorescence imaging during sporulation, showing reporter expression in transformant line #6 compared to WT. Scale bars, 100 µm. **(f)** Detached-leaf infection assay comparing lesion areas produced by WT and MatryoshCaMP8s transformant line #6. Each point represents one inoculation spot. Boxes indicate the interquartile range, center lines indicate medians, and whiskers show the data range. No significant difference was detected between WT and #6 (Kruskal-Wallis test, P = 0.11). n.s.: not significant.

### MatryoshCaMP8s reports rapid cold-shock-induced Ca^2+^ transients in sporangia

We next tested whether MatryoshCaMP8s could report a rapid Ca^2+^-associated physiological response in *Phytophthora*. We focused on cold shock, a treatment commonly used to induce zoospore release and previously shown to trigger Ca^2+^ transients in *Phytophthora cinnamomi* sporangia [30]. Detached *P. palmivora* sporangia were transferred to a custom water-filled microscopy chamber that allowed ice-cold water to be added during live imaging (**Fig. 2a**).

**Figure 2.**
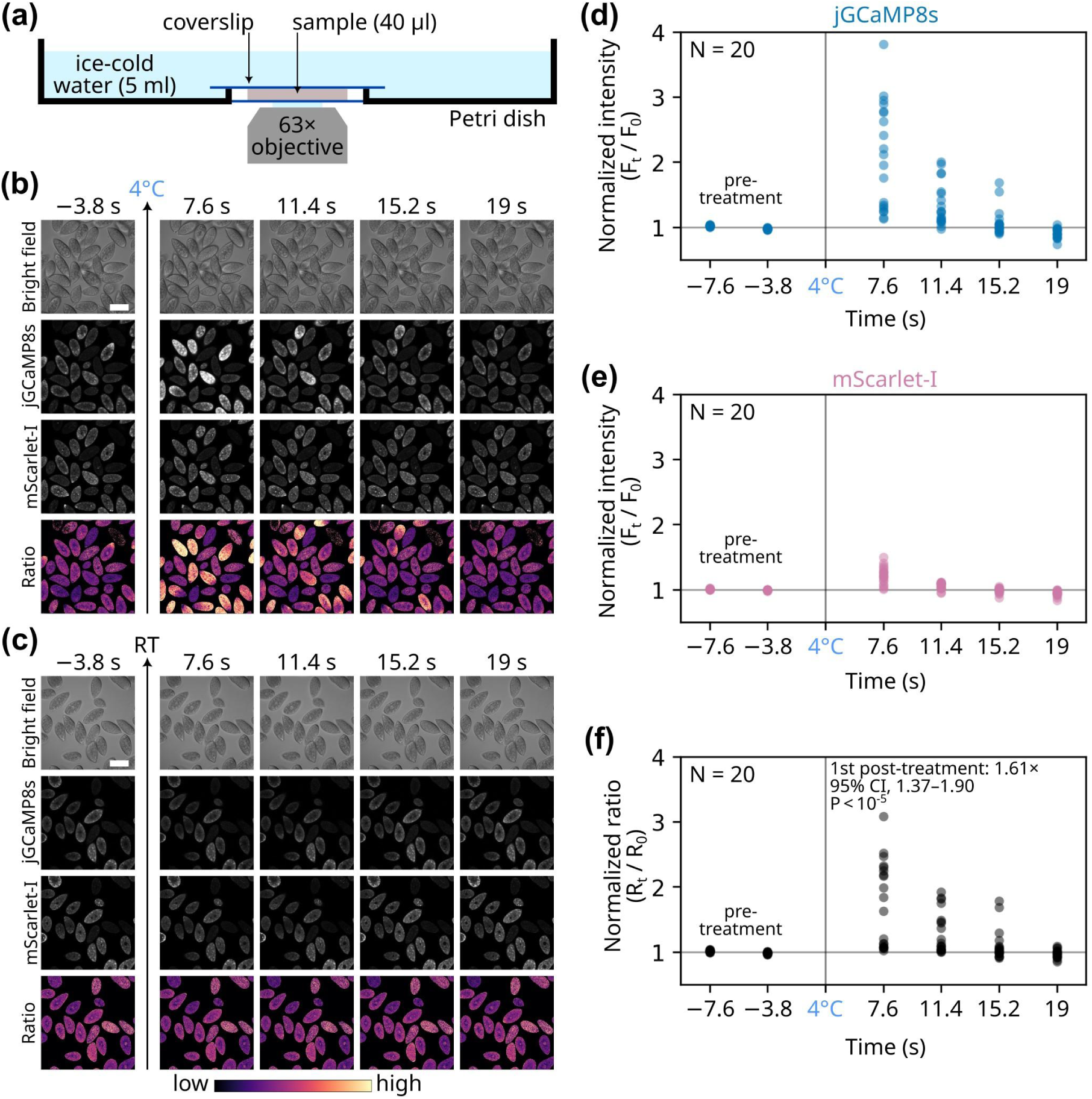
MatryoshCaMP8s reports rapid cold-shock-induced Ca^2+^ responses in *Phytophthora palmivora* sporangia. **(a)** Schematic representation of the custom microscopy setup used to apply cold shock during live imaging. Sporangia were placed in a 40 µl sample confined between two coverslips inside a Petri dish compartment. Ice-cold water (5 ml) was added to the upper compartment after a short pre-treatment phase, used as a reference. **(b)** Representative time series of *P. palmivora* sporangia expressing MatryoshCaMP8s after addition of ice-cold water. Bright field, jGCaMP8s, mScarlet-I, and jGCaMP8s/mScarlet-I ratio images are shown. The vertical arrow indicates the addition of ice-cold water. Frames acquired during focal plane perturbation were excluded. **(c)** Representative control time series acquired after adding room temperature water, using the same protocol. **(d, e)** Quantification of normalized jGCaMP8s fluorescence **(d)** and normalized mScarlet-I fluorescence **(e)** after ice-cold water addition. Each point represents one sporangium. Intensities were normalized to the mean pre-treatment value for each sporangium. **(f)** Quantification of the normalized jGCaMP8s/mScarlet-I ratio after ice-cold water addition. For each sporangium, the ratio was normalized to the mean pre-treatment ratio. At the first post-treatment time point, the mean response reached 1.61-fold relative to pre-treatment values (95% bootstrap CI: 1.37-1.90; one-sample sign-flip permutation test on log_2_-transformed normalized ratios, P < 10^-5^; N = 20 sporangia). The vertical grey line indicates the addition of ice-cold water. Horizontal grey lines indicate the normalized pre-treatment baseline. Scale bars, 50 µm.

jGCaMP8s and mScarlet-I fluorescence were acquired before and after the addition of ice-cold water, using room temperature water as a control (**Fig. 2b, c**). Since water addition transiently perturbed the focal plane, the corresponding frames were excluded from the quantification. Ice-cold water triggered a rapid and transient increase in jGCaMP8s fluorescence in multiple sporangia, whereas the mScarlet-I signal remained comparatively stable (**Fig. 2d, e**). Accordingly, the normalized jGCaMP8s/mScarlet-I ratio increased immediately after cold shock (**Fig. 2f**), reaching 1.61-fold over pre-treatment values at the first post-treatment time point (95% bootstrap CI: 1.37-1.90; P < 10^-5^; N = 20 sporangia). The response returned towards baseline within 20 s, consistent with a fast Ca^2+^ transient and in agreement with earlier Fura-2 dextran measurements [30]. By contrast, adding room temperature water did not induce a comparable increase in jGCaMP8s fluorescence or in the ratio (**Fig. 2c**), indicating that the response was not solely due to water addition or mechanical disturbance. Together, these results provide physiological validation of MatryoshCaMP8s as a reporter of rapid Ca^2+^ signaling events in living *P. palmivora* sporangia.

### Calcimycin immobilizes zoospores and triggers a ratiometric MatryoshCaMP8s response

We further challenged the *P. palmivora* MatryoshCaMP8s line with a pharmacological perturbation expected to increase intracellular Ca^2+^. To this end, we treated swimming zoospores with calcimycin/A23187, a calcium ionophore previously reported to alter zoospore swimming behavior in *Pythium aphanidermatum* [26]. Because ionophore activity can be independently monitored through its effect on zoospore motility, this assay provided a complementary functional test of reporter responsiveness. Time-lapse imaging was performed before and after the addition of 50 µM calcimycin to the medium (**Fig. 3a**). Calcimycin rapidly slowed zoospore movement and triggered encystment (**Fig. 3b**). A similar immobilization phenotype was observed with 100 µM lithium chloride (**Fig. 3b**), consistent with earlier reports linking this treatment to altered intracellular Ca^2+^ homeostasis in *Phytophthora* [34]. For quantification, we identified individual zoospores based on mScarlet-I fluorescence, independently of the jGCaMP8s signal. Calcimycin caused a marked increase in jGCaMP8s fluorescence compared with pre-treatment levels, whereas mScarlet-I fluorescence remained comparatively stable (**Fig. 3c,d,f,g)**. Consequently, the jGCaMP8s/mScarlet-I ratio increased significantly after treatment (**Fig. 3e**). Thus, MatryoshCaMP8s reports pharmacologically induced ratiometric Ca^2+^ responses in living *P. palmivora* zoospores.

**Figure 3.**
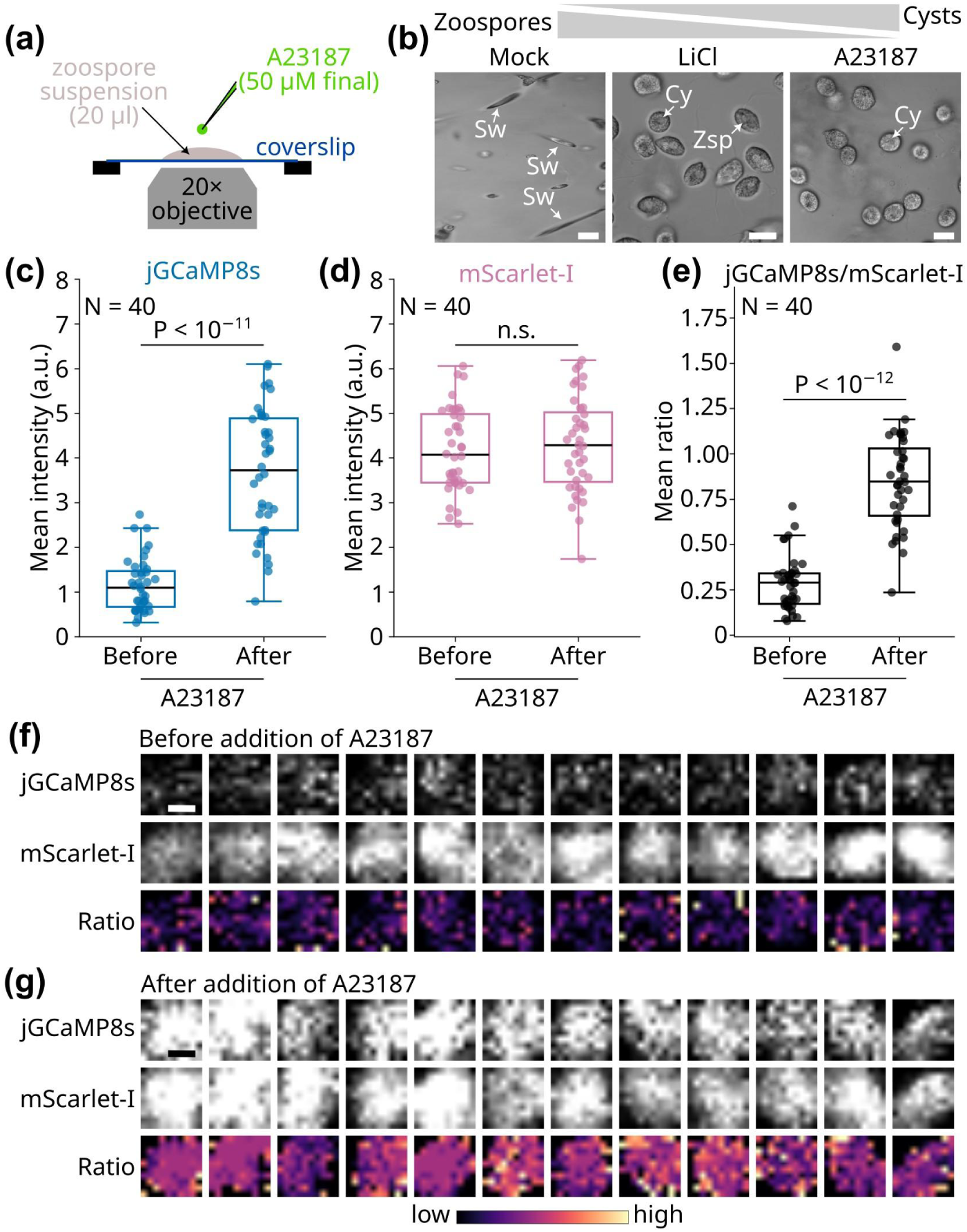
Calcimycin immobilizes zoospores and induces a MatryoshCaMP8s response in *Phytophthora palmivora*. **(a)** Schematic overview of the calcimycin/A23187 treatment assay. A suspension of swimming zoospores was deposited under a coverslip and imaged before and after the addition of calcimycin/A23187 solution (50 µM final). **(b)** Representative bright field images of zoospores after addition of DMSO (Mock), LiCl, or calcimycin/A23187. DMSO-treated zoospores remained motile and are therefore not visible as sharply defined cells in this representative frame, whereas LiCl- and A23187-treated zoospores became immobilized. A23187-treated zoospores rapidly encysted. Scale bars, 10 µm. Sw: swimming zoospore, Zsp: immobile zoospore, Cy: cyst. **(c-d)** Quantification of jGCaMP8s **(c)** and mScarlet-I **(d)** fluorescence intensity in zoospores before and after addition of calcimycin/A23187. Intensities are displayed as mean fluorescence intensity in arbitrary units. Dots indicate individual values (N = 40 distinct zoospores per condition; pre- and post-treatment populations were treated as unpaired samples). P-values were calculated using a two-sided Mann-Whitney test. n.s., not significant. **(e)** Quantification of the jGCaMP8s/mScarlet-I fluorescence ratio before and after calcimycin/A23187 addition. **(f, g)** Representative single-zoospore images showing jGCaMP8s fluorescence, mScarlet-I fluorescence, and the corresponding jGCaMP8s/mScarlet-I ratio before **(f)** and after **(g)** addition of calcimycin/A23187. Scale bars, 5 µm.

### Spatially heterogeneous Ca^2+^ transients precede zoospore release

Having validated MatryoshCaMP8s using both physiological and pharmacological perturbations, we next examined endogenous Ca^2+^ dynamics in sporangia undergoing zoospore differentiation. Detached sporangia were incubated in water and selected based on their granular appearance in bright-field images, indicative of imminent zoospore release (**Fig. 4, Supplementary Movie 1, Zenodo Dataset)**. Time-lapse imaging revealed a stable mScarlet-I signal throughout the acquisition, including immediately before zoospore release. In particular, the mScarlet-I signal was homogeneous within the sporangium, with only occasional puncta (**Fig. 4a**). In contrast, the jGCaMP8s signal exhibited pronounced temporal and spatial dynamics. To quantify these variations, we defined regions of interest (ROIs) within individual sporangia (**Fig. 4b-d**). ROI *a* displayed relatively stable jGCaMP8s intensity and ratio over time (**Fig. 4b**). By contrast, other regions showed transient increases in jGCaMP8s signal and in the corresponding ratio. These dynamics occurred either as successive bursts, as illustrated by ROI *b* (**Fig. 4c**), or as a rapid increase followed by a gradual decrease within ∼30 s, as seen in ROI *c* (**Fig. 4d**). Inspection of the corresponding bright-field images showed minimal movement of the sporangium during these events, making drift or focal-plane changes unlikely explanations for jGCaMP8s signal dynamics. Together, these observations reveal that sporangia are not calcium-homogeneous compartments before zoospore release but instead exhibit localized, transient Ca^2+^ elevations during late zoospore differentiation.

**Figure 4.**
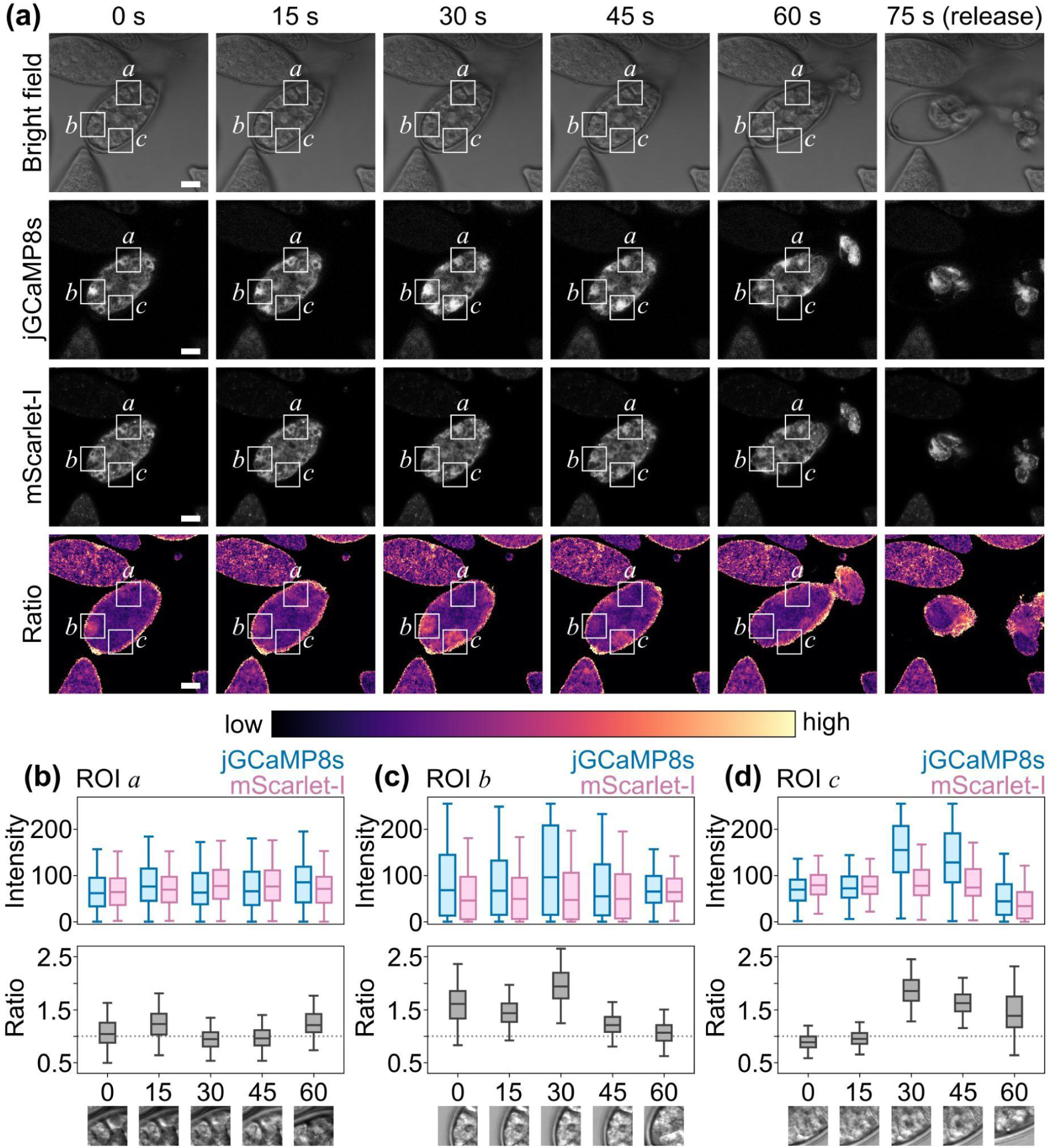
Spatially heterogeneous Ca^2+^ transients precede zoospore release. **(a)** Time-lapse imaging of a representative sporangium undergoing zoospore differentiation and release. Bright field, jGCaMP8s, mScarlet-I, and ratiometric (jGCaMP8s/mScarlet-I) images are shown at the indicated time points. Arrows indicate regions displaying dynamic Ca^2+^ signals. The lookup table represents relative calcium levels (low to high). Scale bars, 10 µm. **(b-d)** Quantification of fluorescence intensity and ratiometric signal in selected regions of interest (ROIs) within the sporangium shown in (a). ROIs (*a*-*c*) correspond to areas indicated in (a). The top panels show jGCaMP8s and mScarlet-I intensities; the bottom panels show the corresponding ratio. Distinct dynamic behaviors are observed, including stable regions (ROI *a*) and transient Ca^2+^ increases occurring as successive bursts (ROI *b*) or as a rapid increase followed by a decrease (ROI *c*). Bright field images corresponding to each time point are shown below the plots for each ROI.

### Rare Ca^2+^ transients occur shortly after zoospore encystment

We next asked whether Ca^2+^ dynamics could be detected shortly after zoospore encystment. Encystment is an essential step for host infection, initiated when zoospores shed their flagella and build a cell wall [35]. To capture Ca^2+^-associated events at this stage, zoospores were mechanically encysted by vortexing, and cysts were imaged immediately by time-lapse confocal microscopy (**Fig. 5, Supplementary Movie 2, Zenodo Dataset)**. mScarlet-I fluorescence gradually decreased in all cysts, consistent with photobleaching of the fluorophore. Although this decrease was partially compensated computationally before ratio analysis, a slight residual decline remained, and late time points were therefore excluded from time-series interpretation (**Fig. 5a**). Most cysts showed stable jGCaMP8s fluorescence during the 5 min acquisition period. However, two out of 45 cysts displayed a transient burst in jGCaMP8s signal, accompanied by corresponding ratiometric increases (**Fig. 5a,b**). In cyst C1, the signal increased over two consecutive frames, lasting ∼20 s in total. In cyst C2, the signal appeared as shorter bursts lasting approximately 10 s each.

**Figure 5.**
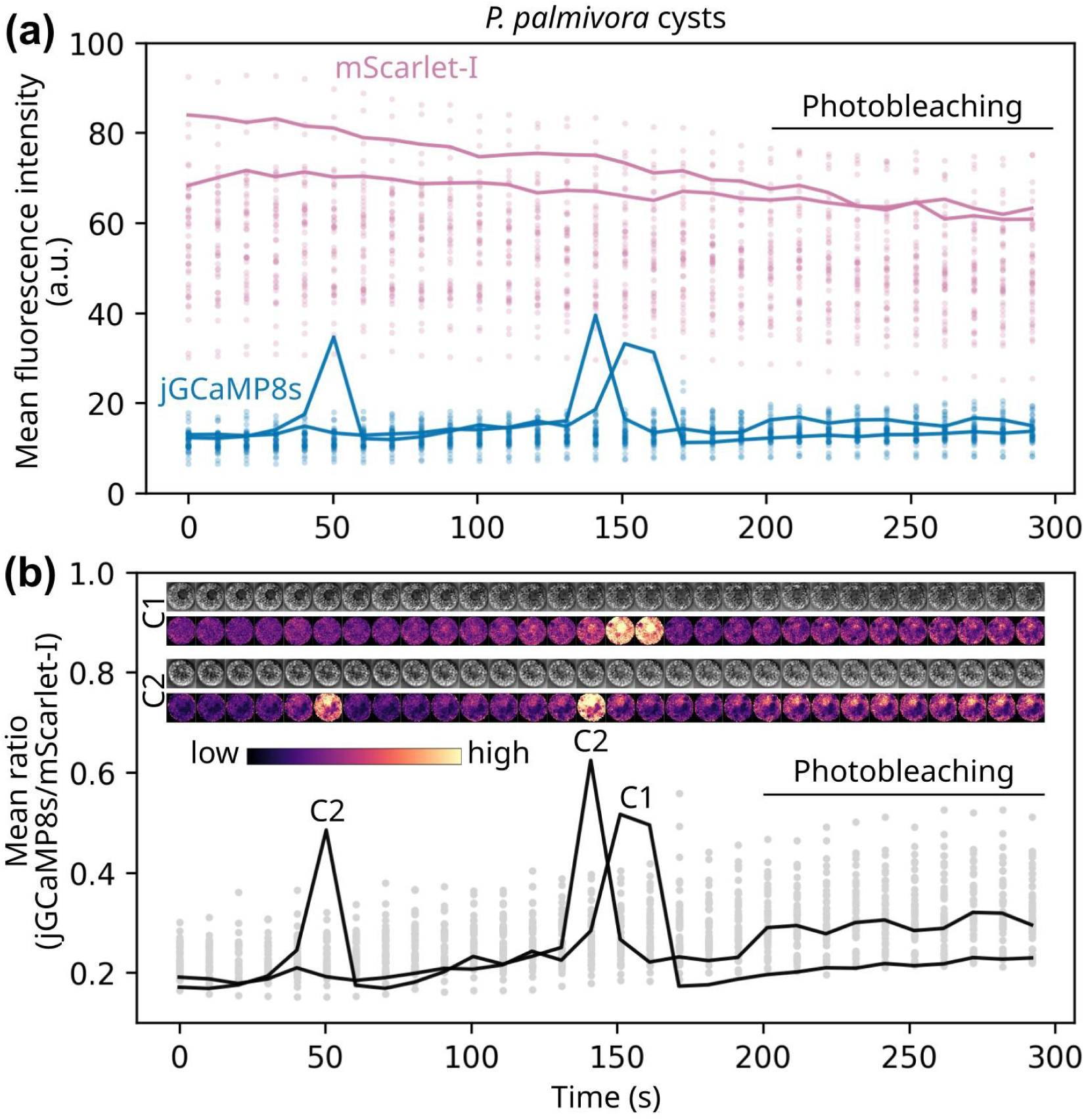
Transient jGCaMP8s fluorescence bursts in *Phytophthora palmivora* cysts. **(a)** Mean fluorescence intensity over time in individual *P. palmivora* cysts expressing the MatryoshCaMP8s reporter. jGCaMP8s and mScarlet-I signals are shown in blue and pink, respectively. Dots represent individual cyst measurements at each time point, while solid lines highlight two representative cysts, C1 and C2, displaying transient increases in jGCaMP8s fluorescence during the acquisition. The gradual decrease in mScarlet-I fluorescence is consistent with photobleaching. a.u.: arbitrary unit. **(b)** Mean jGCaMP8s/mScarlet-I ratio over time for the same cyst population. Grey dots represent individual cyst measurements, and black lines highlight the two representative cysts shown in (a). Image strips show synchronized bright-field and ratio snapshots for C1 and C2 across the time series. Ratio images are displayed using the magma lookup table with identical display scaling across the series; low and high indicate relative ratio values. Late time points are shown for completeness but were not used for biological interpretation.

In both cases, jGCaMP8s fluorescence returned to baseline after the burst, with no corresponding change in the mScarlet-I channel. Bright-field imaging revealed no visible alteration in cyst morphology during acquisition (**Fig. 5b**), indicating that these fluorescence events preceded, or occurred independently of, detectable morphological changes such as germination. Similar bursts were also observed during independent time-lapse acquisitions **(Zenodo Dataset)**, suggesting that these rare events reflect genuine changes in cyst physiology rather than acquisition-specific artifacts. Together, our findings show that MatryoshCaMP8s can reveal rare, transient Ca^2+^ signaling events in cysts that would otherwise remain invisible by morphology alone.

### Ca^2+^ transients occur at the germ tube tip during exploratory growth

We next investigated Ca^2+^ dynamics during cyst germination (**Fig. 6, Zenodo Dataset)**. Swimming zoospores were mechanically encysted by repeated vortexing, and the resulting cysts were incubated on a glass slide in a humid chamber at 25 °C without nutrients. After 1 hour, germinating cysts were imaged by time-lapse confocal microscopy (**Fig. 6a**). We focused on germ tubes displaying a visible terminal swelling in bright-field images, as such swellings have been associated with mechanical forces generated during attempted penetration [9]. Over a two-minute acquisition window, we detected at least four successive peaks of jGCaMP8s fluorescence at the germ tube tip, whereas the mScarlet-I signal remained comparatively stable (**Fig. 6b**). These events coincided with transient increases in the jGCaMP8s/mScarlet-I ratio, supporting their interpretation as Ca²⁺ transients rather than changes in overall fluorescence intensity or focal-plane position. Although our current setup does not allow direct correlation of these Ca^2+^ peaks with discrete biomechanical events at the germ tube tip, their repeated occurrence at the terminal swelling indicates that Ca^2+^ dynamics are active at this prospective penetration site. We next examined whether these tip-associated signals remained local or propagated along the germinating cyst.

**Figure 6.**
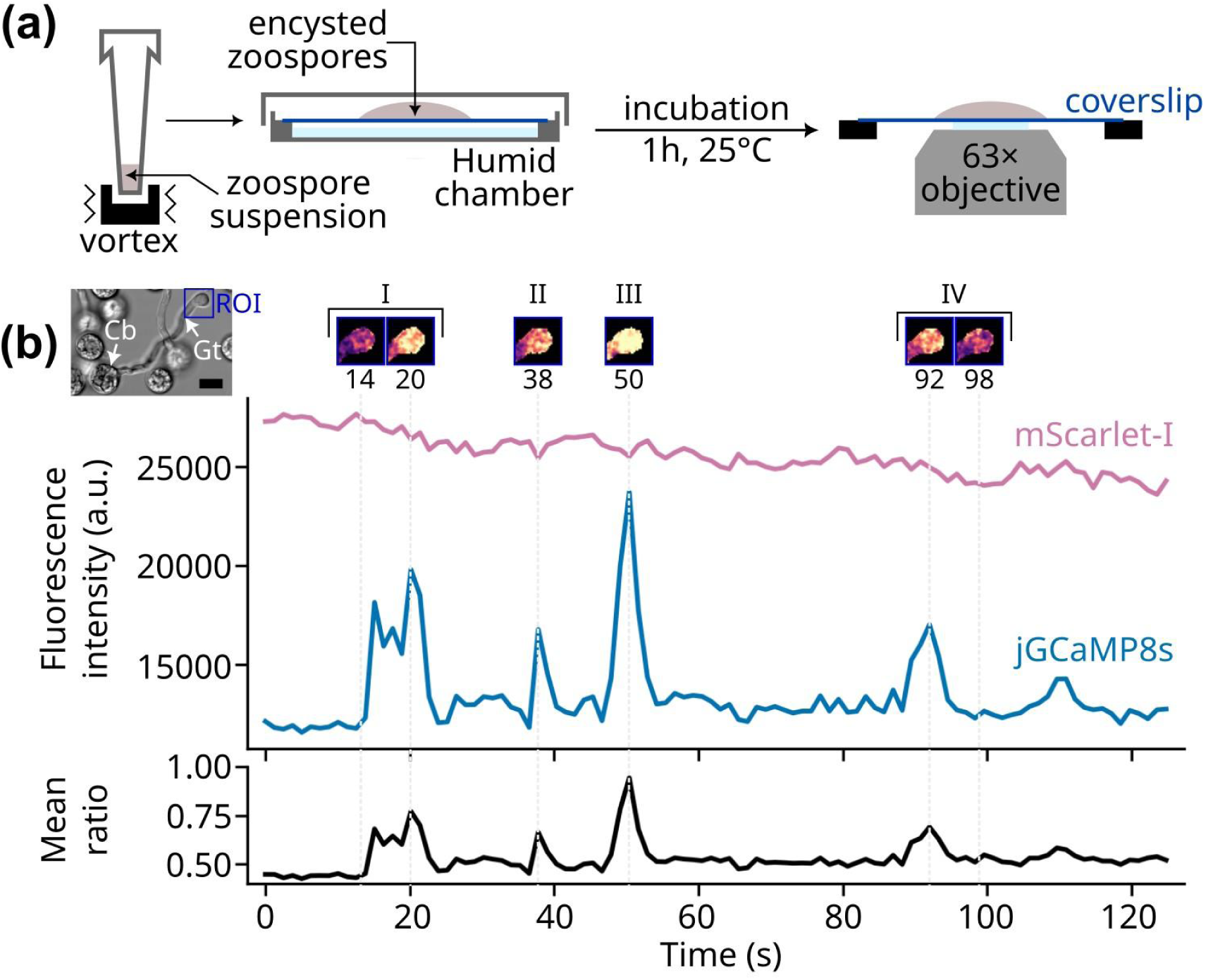
Ca^2+^ transients occur at the germ tube tip during exploratory growth. **(a)** Schematic overview of the experimental setup used to image Ca^2+^ dynamics during cyst germination. Swimming zoospores were mechanically encysted by vortexing, deposited on a glass slide, and incubated in a humid chamber for 1 h at 25°C before time-lapse confocal imaging. **(b)** Representative time-lapse quantification of jGCaMP8s and mScarlet-I fluorescence in a germinating cyst displaying a visible germ tube swelling. Mean fluorescence intensity traces show successive jGCaMP8s peaks (blue) at the germ tube tip, while the mScarlet-I signal (pink) remains comparatively stable. The lower trace shows the corresponding mean jGCaMP8s/mScarlet-I ratio (black). Insets indicate selected time points corresponding to four Ca^2+^ transient events (I-IV). Dashed vertical lines indicate the frames illustrated above the traces. Scale bar, 10 µm.

### Ca^2+^ waves propagate along germ tubes during cyst germination

Germ tubes often grow in three dimensions, making it difficult to track Ca^2+^ dynamics across the entire structure over time. However, in some acquisitions, a germinating cyst remained sufficiently in focus to allow spatially resolved analysis along the germ tube and cyst body. We therefore quantified jGCaMP8s and mScarlet-I fluorescence in sequential regions of interest positioned from the swollen germ tube tip towards the cyst body (ROIs I-XIV; **Fig. 7a, Supplementary Movie 3**). Consistent with the tip-focused analysis, several transient increases in jGCaMP8s fluorescence were detected at the germ tube tip, whereas the mScarlet-I signal remained comparatively stable throughout the acquisition (**Fig. 7b**). These events were mirrored by increases in the jGCaMP8s/mScarlet-I ratio (**Fig. 7c**). Importantly, some Ca^2+^ transients were not confined to the tip but appeared to propagate towards adjacent regions. For example, one event peaked first in tip-proximal ROIs I-II at ∼26.5 s and subsequently reached ROI VII at ∼34.4 s, corresponding to propagation over ∼25 µm in ∼8 s, or an apparent speed of ∼3 µm/s, similar to the speed of some Ca^2+^ waves in certain mammalian cells [36]. Because the germ tube extends in three dimensions and the acquisition rate limits precise wavefront tracking, this value should be interpreted as an approximate propagation speed rather than an absolute measurement. Similar propagating events occurred later in the same germinating cyst, indicating that they were not isolated fluctuations but recurrent spatiotemporal Ca^2+^ dynamics. Together, these observations suggest that germinating cysts can generate Ca²⁺ waves that transmit tip-associated information towards more proximal regions of the germ tube and cyst body, potentially coordinating intracellular organization during early host-colonization-related growth. We next asked whether similar tip-associated Ca^2+^ transients also occur during plant infection.

**Figure 7.**
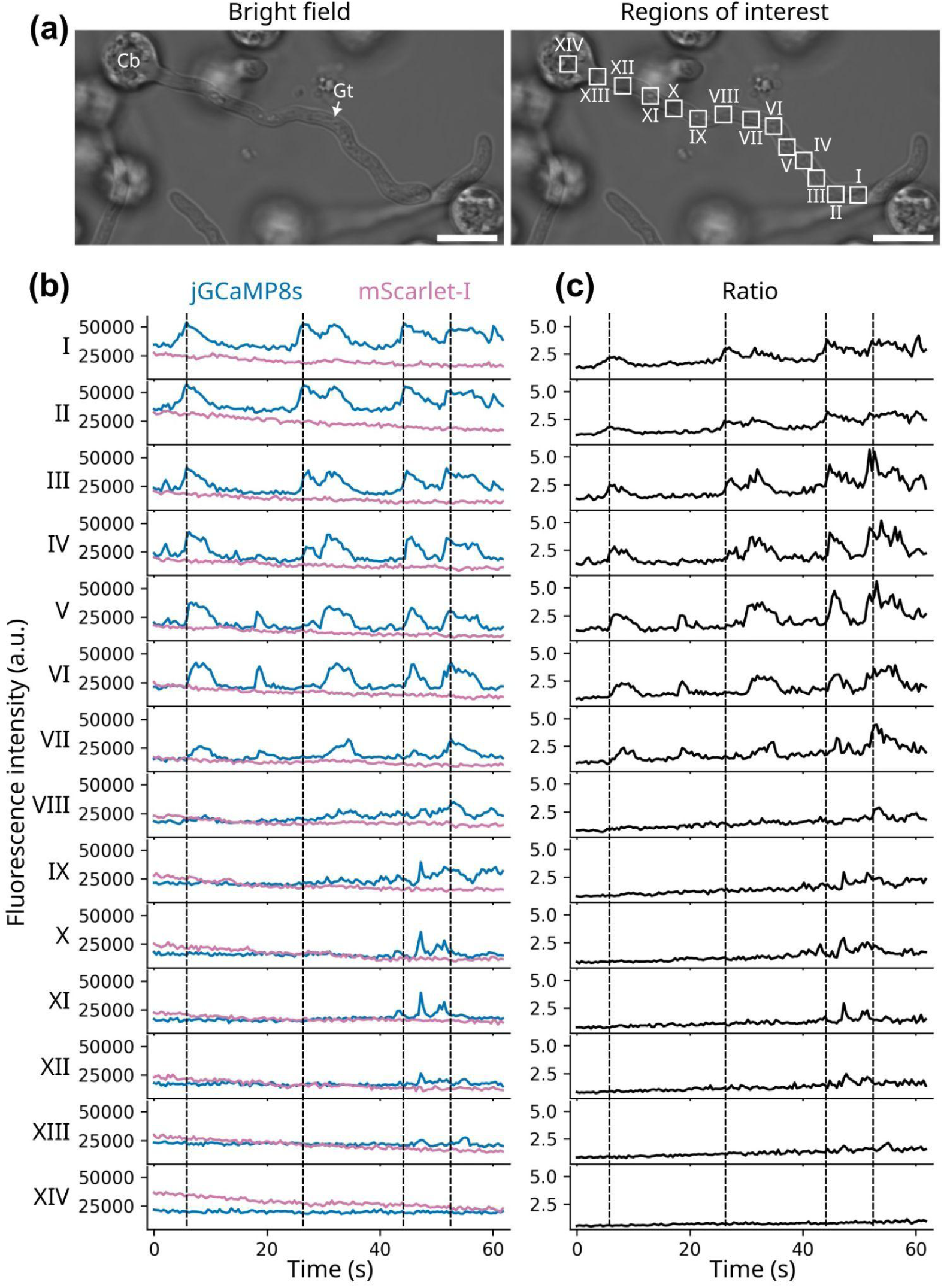
Ca^2+^ waves propagate along germ tubes during cyst germination. **(a)** Bright-field image of a germinating *P. palmivora* cyst and corresponding regions of interest (ROIs) used for spatially resolved fluorescence analysis. ROIs I-XIV were positioned from the swollen germ tube tip towards the cyst body. Cb, cyst body; Gt, germ tube. Scale bars, 10 µm. **(b**) Time-lapse quantification of jGCaMP8s and mScarlet-I fluorescence intensity in ROIs I-XIV. Dashed vertical lines indicate representative Ca^2+^ transient events. **(c)** Corresponding jGCaMP8s/mScarlet-I ratio traces for the same ROIs. Sequential delays in ratiometric peaks across adjacent ROIs indicate propagation of Ca^2+^ signals from the germ tube tip towards more proximal regions of the germ tube and cyst body.

### Tip-associated Ca^2+^ transients occur during early plant infection

After detecting Ca^2+^ transients in cysts germinating *in vitro*, we investigated whether similar dynamics also occur during plant infection. *Nicotiana benthamiana* seedlings were inoculated with *P. palmivora* MatryoshCaMP8s zoospores in a liquid compartment, allowing motile zoospores to reach suitable infection sites along the root surface (**Fig. 8a**). After 3 h of incubation, early infection structures were imaged by time-lapse confocal microscopy. We focused on a germinated cyst displaying a swollen hyphal tip in contact with the root surface, consistent with an attempted penetration event. Shortly after the start of acquisition, jGCaMP8s fluorescence increased sharply, peaking at 6 s before progressively returning towards baseline over the following ∼30 s (**Fig. 8b,c**). Following this initial peak were 5 additional jGCaMP8s bursts at approximately 43, 48, 61, 84, and 90 s (**Fig. 8c**). The most intense of them occurred at 61 s. By contrast, the mScarlet-I signal remained comparatively stable throughout the acquisition. Accordingly, the jGCaMP8s/mScarlet-I ratio displayed two main transient peaks matching the highest increases in jGCaMP8s fluorescence. Towards the end of the acquisition, the morphology of the swollen tip appeared to change slightly. Because active focus stabilization was not used, we cannot exclude a minor contribution of Z-drift to this apparent morphological change. In addition, in the absence of simultaneous biomechanical readout, we cannot determine whether penetration had already begun before the first Ca^2+^ peak or whether the observed pulses preceded this process. These observations show that tip-associated Ca^2+^ transients also occur in cysts germinating at the plant surface, consistent with a link between localized Ca^2+^ signaling and the early stages of host penetration.

**Figure 8.**
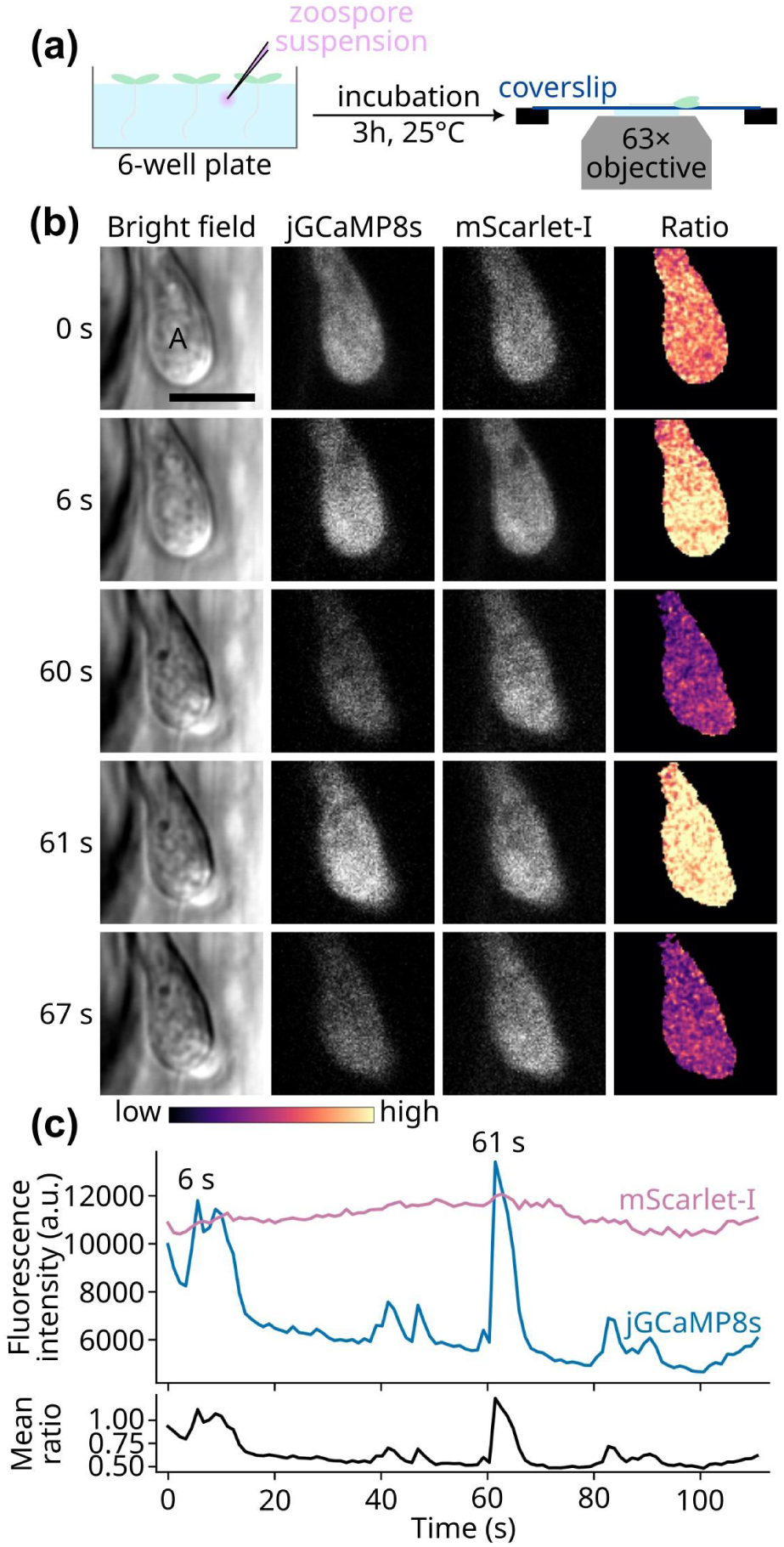
Tip-associated Ca^2+^ transients occur during plant infection. **(a)** Schematic overview of the root infection imaging assay. *Nicotiana benthamiana* seedlings were inoculated with *P. palmivora* MatryoshCaMP8s zoospores and incubated for 3 h at 25 °C before time-lapse confocal imaging. **(b)** Representative time-lapse sequence of an infection structure showing bright-field, jGCaMP8s, mScarlet-I, and jGCaMP8s/mScarlet-I ratio images before, during, and after Ca^2+^ transients. Ratio images are displayed with identical scaling across the entire time series. A, appressorium-like structure. Scale bar, 5 µm. **(c)** Quantification of jGCaMP8s and mScarlet-I fluorescence intensity over time in the infection structure shown in (b). The lower trace shows the corresponding mean jGCaMP8s/mScarlet-I ratio. Two sharp ratiometric Ca^2+^ transients are detected at approximately 6 s and 61 s, while the mScarlet-I signal remains comparatively stable.

## Discussion

Here, we establish live ratiometric calcium imaging in the oomycete pathogen *Phytophthora palmivora* and show that Ca^2+^ signaling during its pre-infective and early infectious development is not a diffuse cellular response, but a series of rapid, spatially restricted events associated with successive developmental and infection-related transitions. By adapting the MatryoshCaMP design to *P. palmivora*, we generated a reporter line suitable for *in vivo* imaging without major detectable effects on sporulation or virulence. We validated MatryoshCaMP8s using two independent perturbations: cold shock, a physiological stimulus previously associated with Ca^2+^ transients in *Phytophthora* sporangia, and calcimycin/A23187, a calcium ionophore that induced both a strong ratiometric reporter response and rapid zoospore immobilization/encystment. We then used this reporter to uncover endogenous Ca^2+^ dynamics across successive stages of the infection cycle, including spatially heterogeneous transients in sporangia prior to zoospore release, rare Ca^2+^ transients shortly after encystment, before any detectable germination, recurrent Ca^2+^ transients at the germ tube tip during exploratory growth, propagating Ca^2+^ waves along germ tubes, and sharp Ca^2+^ pulses during plant infection. Together, these findings reveal stage-specific and subcellular Ca^2+^ dynamics that were previously inaccessible in oomycetes.

Pioneering work by Hardham and colleagues showed that cold shock induces rapid Ca^2+^ elevations in *Phytophthora cinnamomi* sporangia using microinjected Fura-2 dextran [30]. Our observations reproduce this physiological response using a genetically encoded reporter and extend it by resolving subcellular Ca^2+^ events. In particular, we detected localized jGCaMP8s bursts within sporangia approaching zoospore release. Such spatial heterogeneity would have been difficult to capture using microinjection-based measurements. Although the origin of these bursts remains unknown, they may reflect late events in zoospore differentiation and activation. Newly differentiated zoospores remain confined within the sporangial cavity before release, where the onset of motility, repeated contact between neighboring zoospores, or mechanical constraint may trigger local Ca^2+^ entry. This hypothesis is consistent with the presence of candidate mechanosensitive calcium channels in oomycetes, including Piezo-like proteins [8,37]. Testing whether pre-release Ca^2+^ bursts are linked to mechanosensing, flagellar activation, or sporangial discharge will require combining calcium imaging with markers of zoospore motility and controlled mechanical perturbations.

Calcimycin/A23187 provided an additional functional validation of MatryoshCaMP8s in swimming zoospores. The ionophore triggered a marked ratiometric response and rapidly altered zoospore behavior, leading to immobilization and encystment. These observations are consistent with Ca^2+^ elevation being sufficient to perturb zoospore motility and promote encystment-associated transitions, in line with earlier pharmacological studies in oomycetes [26,38]. However, because ionophore treatment imposes a non-physiological Ca^2+^ perturbation, it should be interpreted primarily as evidence that MatryoshCaMP8s reports Ca^2+^-associated changes in living zoospores, and as support for the calcium sensitivity of the zoospore-to-cyst transition, rather than as a reconstruction of the endogenous encystment signal. This distinction is important because the endogenous Ca^2+^ events observed after mechanical encystment were rare, transient, and morphologically silent, suggesting that Ca^2+^ signaling during this stage may be more discrete than pharmacological assays imply.

A major outcome of this study is the identification of recurrent, spatially organized Ca^2+^ dynamics during cyst germination and plant infection. During germination on glass, repeated jGCaMP8s peaks occurred at the germ tube tip while the mScarlet-I reference signal remained comparatively stable, supporting their interpretation as local Ca^2+^ transients rather than imaging artifacts. These events were particularly evident in germ tubes displaying terminal swellings, structures associated with attempted penetration and mechanical force generation [39]. Spatially resolved analysis further showed that some tip-associated Ca^2+^ transients were not strictly local, but propagated from the swollen tip towards more proximal regions of the germ tube and cyst body. This suggests that the germ tube tip may act not only as a site of local Ca^2+^ activity, but also as a source of long-range intracellular Ca^2+^ signals during exploratory growth. Such propagating Ca^2+^ signals may be particularly relevant in coenocytic organisms such as oomycetes, where cytoplasmic continuity requires coordination of cytoskeletal organization and nuclear positioning to maintain an even nuclear distribution across extended hyphal compartments [40]. Importantly, similar sharp ratiometric Ca^2+^ pulses were observed during plant infection, indicating that these dynamics are not restricted to *in vitro* germination conditions but also occur in a host context. This continuity between exploratory growth and infection suggests that polarized Ca^2+^ transients, and in some cases propagating Ca^2+^ waves, are recurrent features of *P. palmivora* development during the transition from environmental exploration to host penetration.

The relationship between Ca^2+^ signaling, mechanical stimulation, and cytoskeletal remodeling is especially relevant in light of recent work on *Phytophthora* appressorium formation. Unlike the melanized appressoria of the rice blast fungus *Magnaporthe oryzae*, *Phytophthora* species form naifu appressoria, in which localized reorganization of the actin cytoskeleton sharpens the invasive hyphal tip to facilitate tissue penetration [9]. Our data do not demonstrate that Ca^2+^ pulses cause actin remodeling. However, the recurrence of sharp Ca^2+^ transients at swollen germ tube tips and infection structures provides a plausible link between local mechanical cues, Ca^2+^ signaling, and cytoskeletal regulation. The propagating Ca^2+^ waves observed along germ tubes further suggest that local tip-associated events may be transmitted to more proximal cellular regions, potentially coordinating cytoskeletal organization, organelle positioning, or nuclear behavior over distances larger than the penetration site itself. Candidate Ca^2+^-responsive cytoskeletal regulators include EF-hand-containing proteins such as α-actinin, as well as actin-nucleating or actin-organizing complexes such as Arp2/3 [10]. The hygromycin-selectable MatryoshCaMP8s line now enables direct testing of these connections by combining calcium imaging with G418-selectable cytoskeletal reporters, including well-established LifeAct-based constructs [41]. Mechanical stimulation assays using deformable substrates, microfluidic confinement, or microneedle indentation could further determine whether local force application is sufficient to trigger Ca^2+^ pulses and downstream cytoskeletal reorganization.

Our work also opens new perspectives for the study of zoospore sensory biology. Zoospore chemotaxis is central to host finding in *Phytophthora* [5]. Recent studies have implicated receptor-like kinases, heterotrimeric G proteins, and Ca^2+^ signaling in this process [42]. However, most Ca^2+^ measurements in oomycetes have relied on chemical dyes or endpoint assays, which are poorly suited to resolve rapid subcellular events [31]. The short duration of the transients observed here suggests that some Ca^2+^-dependent responses may have been missed, blurred, or indirectly inferred in previous studies. In particular, prolonged dye loading in cysts may capture secondary Ca^2+^-associated changes rather than the primary signaling events occurring in motile zoospores. We propose that environmental perception in swimming zoospores may involve rapid Ca^2+^ pulses or waves that reorient motility within seconds. Although real-time confocal imaging of freely swimming zoospores remains technically challenging, the brightness and kinetics of MatryoshCaMP8s, combined with fast acquisition and microfluidic devices, should enable direct analysis of Ca^2+^ dynamics during zoospore navigation, chemotaxis, and host-derived cue perception.

We observed large-amplitude Ca^2+^ transients during pre-infective development and early stages of penetration. While this does not exclude Ca^2+^ signaling during later infection, it suggests that prominent Ca^2+^ pulses associate with polarized growth, mechanical exploration, and early infection structures. The propagation of Ca²⁺ waves from the germ tube tip towards the cyst body also raises the possibility that Ca^2+^ signals coordinate events occurring at a distance from the site of growth, including organelle repositioning or nuclear migration. More localized or compartment-specific Ca²⁺ events may likewise occur at growing hyphal tips, developing haustoria, or sites of nuclear positioning near host interfaces. Future work combining MatryoshCaMP8s with nuclear markers, haustorial reporters, and high-resolution live imaging will help determine whether Ca^2+^ contributes to haustorium biogenesis, intracellular organization, or nuclear behavior during biotrophic growth. By enabling live ratiometric calcium imaging in *P. palmivora*, this work extends dynamic cell biology to oomycetes and other genetically less tractable eukaryotic microbes. To our knowledge, it also provides the first integrated live subcellular view of Ca^2+^ dynamics across the pre-infective developmental cycle and early host colonization in a filamentous plant pathogen. More broadly, it shows how genetically encoded biosensors can uncover rapid, spatially restricted signaling events that would remain hidden in endpoint assays or in comparisons with classical model systems. In *P. palmivora*, this approach reveals polarized Ca^2+^ transients as recurrent features of developmental transitions and early host colonization, placing Ca^2+^ dynamics at the interface between motility, polarity, mechanics, and infection.

## Materials and methods

### Microbial strains and culture conditions

*Phytophthora palmivora* Butler isolate LILI (accession no. P16830) was originally isolated from oil palm in Colombia and is maintained in the *Phytophthora* collection at the Sophia Agrobiotech Institute (Sophia Antipolis, France). *P. palmivora* strains were cultured on V8 agar plates (1.5% agar) at 25°C under constant light conditions. Transformed strains were maintained in the presence of hygromycin at a final concentration of 50 mg/L. For zoospore production, 7-day-old plates were incubated at 4°C for 30 min, then flooded with sterile water and incubated at room temperature for 15 min.

### Plant material and growth conditions

*Nicotiana benthamiana* plants were grown at 24°C for 4 weeks under a 16h light/8h dark cycle. Plants were cultivated in the presence of biocontrol agents *Steinernema feltiae* (Koppert, The Netherlands) and *Bacillus thuringiensis* (Valent BioSciences, USA). For *in vitro* assays, *N. benthamiana* seedlings were grown for one week as previously described [43].

### Plasmid construction

The MatryoshCaMP8s construct was based on the design described by [24]. jGCaMP8s, designed and optimized by [22], was used as the backbone. An mScarlet-I carrying the E95D substitution [33], corresponding to E220D in the full MatryoshCaMP8s reporter, was inserted into a disordered region of jGCaMP8s and flanked by GGT/GGS linkers. jGCaMP8s and the mutated mScarlet-I sequences were inserted by overlap extension PCR into a modified pGEMT plasmid containing a hygromycin resistance cassette. Plasmid assembly was performed using NEBuilder HiFi DNA Assembly (New England Biolabs, USA).

### MatryoshCaMP8s protein structure

The MatryoshCaMP8s structure was predicted using AlphaFold 3 running on AlphaFold server (https://alphafoldserver.com/). The protein model of the biosensor was rendered and analyzed with UCSF ChimeraX (https://www.cgl.ucsf.edu/chimerax/).

### *Phytophthora* transformation

*P. palmivora* was transformed by electroporation. The electroporation mixture contained 80 µg of plasmid DNA, modified Petri’s solution (final concentrations: 0.25 mM CaCl_2_, 1 mM MgSO_4_, 1 mM KH_2_PO_4_, and 0.8 mM KCl), and zoospores, brought to a final volume of 800 µL. Electroporation was performed using a Gene Pulser Xcell Electroporation System (Bio-Rad, USA) with exponential decay, 500 V, 50 µF, and 800 ohms. After electroporation, zoospores were incubated for 8 h at 25°C in liquid V8 medium with gentle shaking. Transformants were selected on V8 agar plates containing hygromycin (50 mg/L).

### Cyst assays

Cysts were obtained by mechanically inducing encystment of freshly released zoospores through vortexing as previously described [44]. Cysts were then transferred onto a glass coverslip and immediately imaged. For cyst germination assays, coverslips were placed in a humid chamber and incubated for 1 h prior to imaging.

### Infection assays

Detached leaves and root inoculations were performed using the *P. palmivora* wild-type LILI strain and transgenic line #6. Middle leaves (P2) were used for detached leaf infection assays. Leaves were inoculated with droplets containing 1000 zoospores and incubated for 2 days at 25°C under constant light in a humid environment. After incubation, leaf images were acquired with the IDS Cockpit software (IDS, Germany) using a blue-light transilluminator (Thermo Fisher Scientific, USA) and subsequently analyzed with ImageJ (https://imagej.net/ij/). Root inoculations were performed on 1-week-old *N. benthamiana* seedlings. Seedlings were transferred to sterile water, after which zoospore suspensions were added at varying concentrations to optimize infection density while maintaining imaging accessibility. Inoculated roots were observed after 3 hours of incubation.

### Confocal microscopy

Microscopic observations were performed using an LSM 880 confocal laser-scanning microscope (Zeiss, Germany) equipped with a 63×, NA 1.2, water-immersion objective. The cpGFP of jGCaMP8s and the mutated Scarlet-I fluorophores were excited at 488 and 561 nm, respectively.

For time-lapse imaging of immobile objects, mScarlet-I photobleaching was compensated for by rescaling each frame to match the initial reference intensity while preserving background levels [45]. A reference signal was computed within a mask defined from the initial frame and consistently applied across time, under the assumption that the selected region remains structurally stable. Briefly, at each time point *t*, a background-corrected signal *S_t_* was computed, and a correction factor *f_t_* was defined relative to the initial frame as:

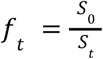

The corrected intensity *F^corr^_r_* was then obtained by:

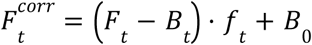

where *F_t_*is the raw intensity, *B_t_* the background at time *t*, and *B*_0_ the background at the initial time point.

### Ratiometric image analysis

For ratiometric analyses, pixel-wise ratios were computed between the numerator (cpGFP) and the denominator (mScarlet-I) channels. For each pixel, the raw ratio *R* was calculated as:

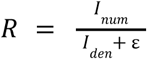

where *I_num_* and *I_den_* are the 16-bit intensities of the numerator and denominator channels, respectively, and ε is a small constant introduced to avoid division by zero (ε = 10⁻⁶). Ratios were computed only within a valid mask defined by signal intensity thresholds and morphological filtering. These raw ratio values were saved for quantitative analysis without further transformation.

For visualization purposes, ratio images were independently rescaled using percentile-based clipping (typically 2nd-98th percentiles) and mapped to the “magma” lookup table using the Matplotlib library (https://matplotlib.org/). Depending on the analysis, scaling was applied per frame, per time series, or across a full dataset to ensure consistency. Rescaling was applied solely for display and did not affect the raw ratio values used for downstream quantification.

### Statistical analysis

For cold shock response assays, the jGCaMP8s/mScarlet-I fluorescence ratio R was calculated for each sporangium and normalized to the mean pre-treatment ratio R_0_. Normalized ratios were log_2_-transformed before statistical analysis, so that no change corresponded to 0, and reciprocal fold changes were treated symmetrically. The primary response metric was defined as the log_2_(R/R_0_) value at the first post-treatment time point. As secondary descriptive metrics, we also computed the peak post-treatment log_2_(R/R_0_) value and the integrated post-treatment response, quantified as the area under the log_2_(R/R_0_) curve using the trapezoidal rule. Statistical significance was assessed using a one-sample sign-flip permutation test against zero, with sporangia as the experimental units. Confidence intervals for mean responses were estimated by bootstrap resampling of sporangia.

## Data and code availability

Time-lapse videos corresponding to the figures are provided as Supplementary Movies. Additional representative time-lapse acquisitions not used for figure generation are available on Zenodo at https://doi.org/10.5281/zenodo.20137233. Raw imaging data and other source data are available from the corresponding author upon reasonable request. Scripts used for image processing, ratiometric analysis, and figure generation are available on GitHub at https://github.com/EEvangelisti/MatryoshCaMP8s.

## Acknowledgements

We thank Sander Schouten (WUR) and Sebastian Schornack (SLCU) for insightful discussions during the early stages of the project, and Francine Govers (WUR) and Tijs Ketelaar (WUR) for valuable feedback. We acknowledge the PlantBios Imaging Facility at the Sophia Agrobiotech Institute for technical assistance with confocal microscopy, and Agnès Attard (ISA) for advice on confocal imaging assays.

## Author contributions

M.H.J.P.: investigation, formal analysis, visualization. D.A.: investigation. J.J.M.: conceptualization and experimental design. E.E.: conceptualization, funding acquisition, supervision, formal analysis, visualization, and writing, original draft. All authors reviewed and approved the manuscript.

## Supplementary Movies

**Supplementary Movie 1.** Live imaging of Ca^2+^ dynamics during *Phytophthora palmivora* sporangial discharge.

**Supplementary Movie 2.** Live imaging of Ca^2+^ dynamics in newly formed *Phytophthora palmivora* cysts.

**Supplementary Movie 3.** Live imaging of Ca^2+^ dynamics in *Phytophthora palmivora* cysts germinating on glass.

**An extended supporting dataset** is available on Zenodo at https://doi.org/10.5281/zenodo.20137233.

## References

1. Abad ZG, Burgess TI, Bourret T, Bensch K, Cacciola SO, Scanu B, et al. *Phytophthora*: taxonomic and phylogenetic revision of the genus. Stud Mycol. 2023;106: 259–348.

2. Coomber A, Saville A, Ristaino JB. Evolution of *Phytophthora infestans* on its potato host since the Irish potato famine. Nat Commun. 2024;15: 6488.

3. Wang W-J, Li J, Zhao G, Wang Y-P, Fan S, Dong Y-Q, et al. Pathogenicity and virulence of *Phytophthora infestans*: The ever-evolving threat to food security and its sustainable management strategies. Virulence. 2025;16: 2586882.

4. Torres GA, Sarria GA, Martinez G, Varon F, Drenth A, Guest DI. Bud rot caused by *Phytophthora palmivora*: a destructive emerging disease of oil palm. Phytopathology. 2016;106: 320–329.

5. Kasteel M, Ketelaar T, Govers F. Fatal attraction: How *Phytophthora* zoospores find their host. Semin Cell Dev Biol. 2023; 148–149: 13–21.

6. Judelson HS, Blanco FA. The spores of *Phytophthora*: weapons of the plant destroyer. Nat Rev Microbiol. 2005;3: 47–58.

7. Wang Y, Govers F, Wang Y. Oomycete plant pathogens: biology, pathogenesis and emerging control strategies. Nat Rev Microbiol. 2026;24: 270–287.

8. Evangelisti E, Govers F. Roadmap to success: How Oomycete plant pathogens invade tissues and deliver effectors. Annu Rev Microbiol. 2024;78:493–512.

9. Bronkhorst J, Kasteel M, van Veen S, Clough JM, Kots K, Buijs J, et al. A slicing mechanism facilitates host entry by plant-pathogenic *Phytophthora*. Nat Microbiol. 2021;6: 1000–1006.

10. Bronkhorst J, Kots K, de Jong D, Kasteel M, van Boxmeer T, Joemmanbaks T, et al. An actin mechanostat ensures hyphal tip sharpness in *Phytophthora infestans* to achieve host penetration. Sci Adv. 2022;8: eabo0875.

11. Romito O, Guéguinou M, Raoul W, Champion O, Robert A, Trebak M, et al. Calcium signaling: A therapeutic target to overcome resistance to therapies in cancer. Cell Calcium. 2022;108: 102673.

12. Luan S. Calcium signaling in plants: Universal and unique paradigms. Cell. 2026;189: 1001–1023.

13. Eisner D, Neher E, Taschenberger H, Smith G. Physiology of intracellular calcium buffering. Physiol Rev. 2023;103: 2767–2845.

14. Park C-J, Shin R. Calcium channels and transporters: Roles in response to biotic and abiotic stresses. Front Plant Sci. 2022;13: 964059.

15. Enomoto M, Nishikawa T, Siddiqui N, Chung S, Ikura M, Stathopulos PB. From stores to sinks: Structural mechanisms of cytosolic calcium regulation. Krebs J, editor. Adv Exp Med Biol. 2017;981: 215–251.

16. Jiang Y, Ding P. Calcium signaling in plant immunity: a spatiotemporally controlled symphony. Trends Plant Sci. 2023;28: 74–89.

17. Lai R, Li G, Cui Q. Flexibility of binding site is essential to the Ca2+ selectivity in EF-hand calcium-binding proteins. J Am Chem Soc. 2024;146: 7628–7639.

18. Negi NP, Prakash G, Narwal P, Panwar R, Kumar D, Chaudhry B, et al. The calcium connection: exploring the intricacies of calcium signaling in plant-microbe interactions. Front Plant Sci. 2023;14: 1248648.

19. Oldroyd GED, Downie JA. Nuclear calcium changes at the core of symbiosis signalling. Curr Opin Plant Biol. 2006;9: 351–357.

20. Ghosh S, Dahiya M, Kumar A, Bheri M, Pandey GK. Calcium imaging: a technique to monitor calcium dynamics in biological systems. Physiol Mol Biol Plants. 2023;29: 1777–1811.

21. Contini C, Kuntz J, Massing U, Merfort I, Winkler K, Pütz G. On the validity of fluorimetric intracellular calcium detection: Impact of lipid components. Biochem Biophys Res Commun. 2023;643: 186–191.

22. Zhang Y, Rózsa M, Liang Y, Bushey D, Wei Z, Zheng J, et al. Fast and sensitive GCaMP calcium indicators for imaging neural populations. Nature. 2023;615: 884–891.

23. Kim H-S, Czymmek KJ, Patel A, Modla S, Nohe A, Duncan R, et al. Expression of the Cameleon calcium biosensor in fungi reveals distinct Ca(2+) signatures associated with polarized growth, development, and pathogenesis. Fungal Genet Biol. 2012;49: 589–601.

24. Ast C, Foret J, Oltrogge LM, De Michele R, Kleist TJ, Ho C-H, et al. Ratiometric Matryoshka biosensors from a nested cassette of green- and orange-emitting fluorescent proteins. Nat Commun. 2017;8: 431.

25. von Broembsen SL, Deacon JW. Calcium interference with zoospore biology and infectivity of Phytophthora parasitica in nutrient irrigation solutions. Phytopathology. 1997;87: 522–528.

26. Donaldson SP, Deacon JW. Changes in motility of *Pythium* zoospores induced by calcium and calcium-modulating drugs. Mycol Res. 1993;97: 877–883.

27. Connolly MS, Williams N, Heckman CA, Morris PF. Soybean isoflavones trigger a calcium influx in *Phytophthora sojae*. Fungal Genet Biol. 1999;28: 6–11.

28. Hyde GJ, Heath IB. Ca^2+^ gradients in hyphae and branches of *Saprolegnia ferax*. Fungal Genet Biol. 1997;21: 238–251.

29. Lew RR. Comparative analysis of Ca^2+^ and H^+^ flux magnitude and location along growing hyphae of *Saprolegnia ferax* and *Neurospora crassa*. Eur J Cell Biol. 1999;78: 892–902.

30. Jackson SL, Hardham AR. A transient rise in cytoplasmic free calcium is required to induce cytokinesis in zoosporangia of *Phytophthora cinnamomi*. Eur J Cell Biol. 1996;69: 180–188.

31. Pei Y, Ji P, Miao J, Gu X, Wang H, Zhao Y, et al. A receptor kinase senses sterol by coupling with elicitins in auxotrophic *Phytophthora*. Proc Natl Acad Sci USA. 2024;121. doi:10.1073/pnas.2408186121

32. Ross BL, Tenner B, Markwardt ML, Zviman A, Shi G, Kerr JP, et al. Single-color, ratiometric biosensors for detecting signaling activities in live cells. Elife. 2018;7. doi:10.7554/eLife.35458

33. Fenno LE, Ramakrishnan C, Kim YS, Evans KE, Lo M, Vesuna S, et al. Comprehensive dual- and triple-feature intersectional single-vector delivery of diverse functional payloads to cells of behaving mammals. Neuron. 2020;107: 836–853.e11.

34. Érsek T, Hölker U, Höfer M. Non-lethal immobilization of zoospores of *Phytophthora infestans* by Li+. Mycol Res. 1991;95: 970–972.

35. Mitchell HJ, Kovac KA, Hardham AR. Characterisation of *Phytophthora nicotianae* zoospore and cyst membrane proteins. Mycological Research. 2002;106: 1211–1223.

36. Fazekas F, Vasbányai L, Berekméri E. Intracellular Ca^2+^ waves in mammalian cells. Biol Futur. 2025;76: 293–313.

37. Zheng L, Mackrill JJ. Calcium signaling in Oomycetes: An evolutionary perspective. Front Physiol. 2016;7: 123.

38. Hyde GJ, Heath IB. Ca2+-dependent polarization of axis establishment in the tip-growing organism, Saprolegnia ferax, by gradients of the ionophore A23187. Eur J Cell Biol. 1995;67: 356–362.

39. Hardham AR. Cell biology of plant-oomycete interactions. Cell Microbiol. 2007;9: 31–39.

40. Evangelisti E, Shenhav L, Yunusov T, Le Naour-Vernet M, Rink P, Schornack S. Hydrodynamic shape changes underpin nuclear rerouting in branched hyphae of an oomycete pathogen. mBio. 2019;10: e01516–19.

41. Kots K, Meijer HJG, Bouwmeester K, Govers F, Ketelaar T. Filamentous actin accumulates during plant cell penetration and cell wall plug formation in *Phytophthora infestans*. Cell Mol Life Sci. 2017;74: 909–920.

42. Ji P, Bao Y, Zhou H, Pei Y, Song W, Ou K, et al. Blocking the isoflavone chemoreceptor in *Phytophthora sojae* to prevent disease. Sci Adv. 2025;11:eadt0925.

43. Evangelisti E, Gogleva A, Hainaux T, Doumane M, Tulin F, Quan C, et al. Time-resolved dual transcriptomics reveal early induced *Nicotiana benthamiana* root genes and conserved infection-promoting *Phytophthora palmivora* effectors. BMC Biol. 2017;15: 39.

44. Lupatelli CA, Seassau A, Magliano M, Kuhn ML, Rey A, Poët M, et al. Membrane proteome analysis identifies key components of sensing in *Phytophthora parasitica* zoospores. Sci Rep. 2025;15:23500.

45. Miura K. Bleach correction ImageJ plugin for compensating the photobleaching of time-lapse sequences. F1000Res. 2020;9: 1494.

